# Rapid Rerouting of Myosin Traffic at the T Cell Immunological Synapse

**DOI:** 10.1101/2022.04.06.487340

**Authors:** Robert Mąka, Natalia Plewa, Urszula Cichoń, Katarzyna Krysztofiak, Jagoda J. Rokicka, Ronald S. Rock

**Author notes:** Email addresses:* (Robert Mąka), (Natalia Plewa), (Urszula Cichoń), (Katarzyna Krysztofiak), (Jagoda J. Rokicka), (Ronald S. Rock). Equal contribution (Robert Mąka, Natalia Plewa).

## Abstract

Cytoskeletal motors travel in patterns set by the architecture of their tracks. Nevertheless, we have a limited understanding of how cells dynamically reorganize their traffic patterns in response to signaling events. To investigate cytoskeletal motor rerouting, we used T cells as a model system. Upon an encounter between a T cell and an antigen presenting cell, the T cell builds a specialized interface with spatially organized immunoreceptors and adhesion molecules called the immunological synapse (IS). The IS also constructs new actin networks within minutes that define the synaptic structure. Here we track the movements of single myosin motors along presynaptic and synaptic actin networks of the T cell. We find that both myosin-5 and myosin-6 reroute after IS construction. For example, most myosin-5 traffic moves inward at the IS, although most of the IS actin filaments have a barbed end out orientation. This anomalous myosin-5 traffic pattern indicates that the IS makes two types of actin networks: a structural network that controls IS shape, and a distinct trafficking network that supports myosin motility. We disrupt these trafficking networks with chemical probes against actin, which inhibits the appearance of cell surface markers of T cell activation. Our results highlight the importance of the sparse actin networks at the center of the IS in T cell function.

## Introduction

T cells form specialized contacts with other cells, which is essential for their maturation and effector function. For example, T cells activate when they contact antigen presenting cells (APCs) displaying antigens in the early stages of the adaptive immune response. One of the early events in T cell activation is the construction of an immunological synapse (IS) with the APC [23, 43]. This synapse focuses signaling between the two cells and helps ensure the fidelity of T cell activation. The IS is a prominent actin-based structure of concentric rings that the T cell constructs within minutes [46, 31]. After activation, T cells undergo clonal expansion and differentiation, and ultimately carry out effector functions throughout the body.

Activated T cells also create synapses as part of their effector function. Cytotoxic CD8^+^ T cells scan tissues searching for cells that display viral antigens on their major histocompatibility complexes (MHCs). When the T cell encounters this virus-infected cell, it forms a synapse to focus secretion of cytotoxic components (granzymes and perforins) to the infected cell. Likewise, helper CD4^+^ T cells form synapses to secrete cytokines for paracrine regulation of other immune cells [17]. Thus, the synapse is a structure of central importance in immune system function. Moreover, the T cell may tune features of its IS structure to support specific effector functions [42].

The IS structure allows the T cell to form a seal with and to focus its responses toward a specific cell. In general, the IS forms three zones in a series of concentric rings called the distal-, proximal-, and central supramolecular activation clusters (dSMAC, pSMAC, and cSMAC) [46]. Each zone has its own set of cell surface markers and specialized actin filament networks [10, 26]. Construction of the IS requires careful coordination of actin polymerization and myosin motor activity [19, 47, 25, 29].

Because the IS is a structure with rich and functionally important cytoskeletal dynamics, we wondered how myosin transport systems evolve over the course of IS formation. The role of nonmuscle myosin-2A in establishing the structure of the dSMAC and pSMAC is by now well understood [47, 29]. However, the activity of transport myosins such as myosin-5 and myosin-6 at the IS is unknown.

To answer questions such as these, we have developed a motility assay that allows us to track myosin movements along cellular actin filaments. Much like the original *Nitella* motility assay [38], we open cells and expose their actin to labeled myosins for tracking. We call this the *ex vivo* motility assay to refer to the source of the actin filaments. Several groups have used this *ex vivo* motility approach to study various systems [7, 41, 8, 20]. By tracking single myosins along actin from single cells, we learn how the cell patterns its actin and how the myosin responds.

Here, we apply this functional imaging approach to show how T cells remodel their myosin traffic patterns along with their actin networks at the immunological synapse. We find that myosins reroute their traffic after T cell activation: myosin-5 switches from centrifugal to centripetal movement, while myosin-6 switches from centripetal to circulatory movement. Small molecule inhibitors of actin network structure and function also perturb the traffic of these myosins. One inhibitor in particular affects the display of cell surface markers of T cell activation. Together, we find that T cells regulate and redirect myosin traffic during IS construction and T cell activation.

## Results

### Myosin-5 travels outward in resting T cells but inward at the immunological synapse

To image the actin networks and myosin traffic at the mature immunological synapse, we use a common approach with antibody-coated coverslips to begin synapse formation in the image plane [24, 28, 47]. One antibody binds and activates through CD3, while a second antibody binds CD28 to provide a co-stimulatory interaction (mouse anti-human OKT3 and CD28.2 antibodies, respectively). We omit the anti-CD3 antibody in our non-activating control condition. We apply T cells to these coverslips, allow cells to adhere or synapses to form, and then fix the actin in place and remove the plasma membrane.

We then add labeled myosin-5 or myosin-6 for tracking by total internal reflection microscopy (Fig. 1). Under these conditions, we detect hundreds to thousands of runs per cell over several minutes. Our complete dataset includes recordings from 511 cells and 340,000 myosin runs, distributed over activation surfaces, cell type, and cytoskeletal inhibitor treatments (Supp. Table S1).

**Figure 1:**
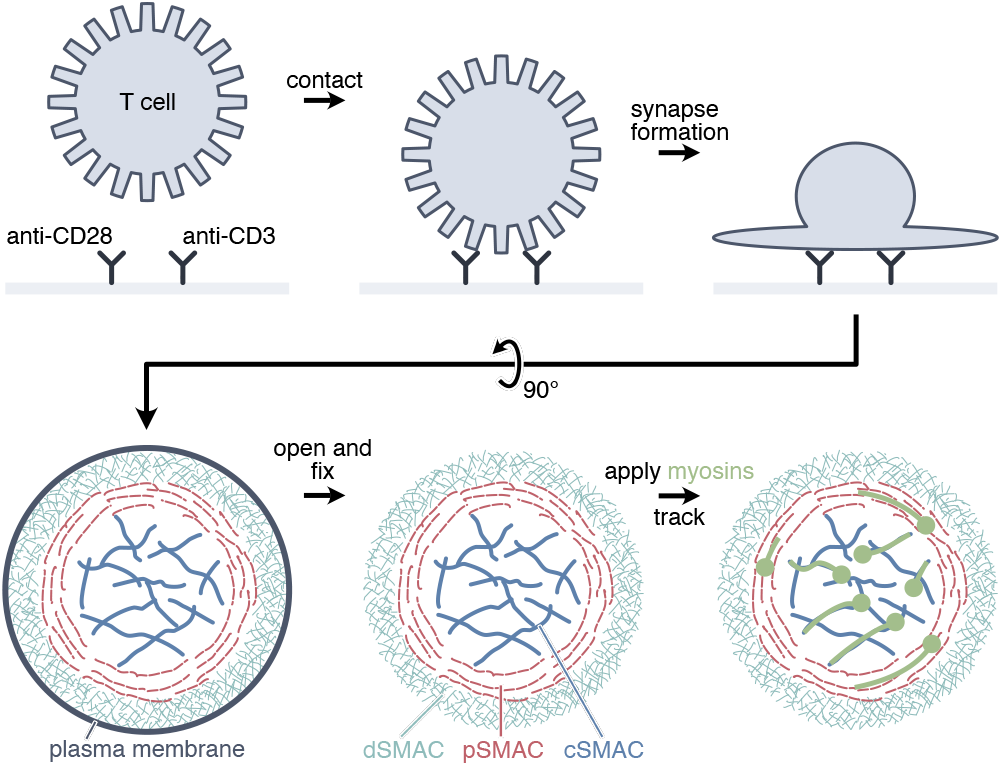
The *ex vivo* motility assay scheme applied to the immunological synapse. We apply lymphocytes (Jurkat T cells, human PBMCs, or CD8^+^ T cells from PBMCs) to a coverslip coated with activating antibodies. After contact with an activating surface, an immunological synapse forms within 5 min. We then open the cell with detergent and stabilize and label the actin filaments to visualize the synapse. Finally, we apply fluorophore-labeled myosin motors (myosin-5 or myosin-6) and track their movements through the synapse. Perturbations include activating vs. non-activating surfaces (anti-CD3/CD28 vs anti-CD28 alone) and small molecule inhibitors of actin filament assembly and function (CK666 and TR100).

On non-activating anti-CD28 surfaces, Jurkat T cells adhere but do not remodel their actin networks to form synapses. Examples of these resting cells and their myosin-5 runs appear in Fig. 2A, top, and Video S1. To show where myosin-5 commences runs we calculate the myosin landing density, a kernel density estimate obtained from the starting points of all runs. Although myosin-5 travels throughout the cell, it concentrates on surface-attached microvilli projecting out from the edge of the cell. Myosin-5 travels out, away from the cell center and toward the tips of the microvilli, as shown in the colored traces shown in the second row of Fig. 2A. The barbed-end-out polarity of the actin core of the microvillus sets this outward movement, as myosin-5 is a barbed-end directed myosin.

**Figure 2:**
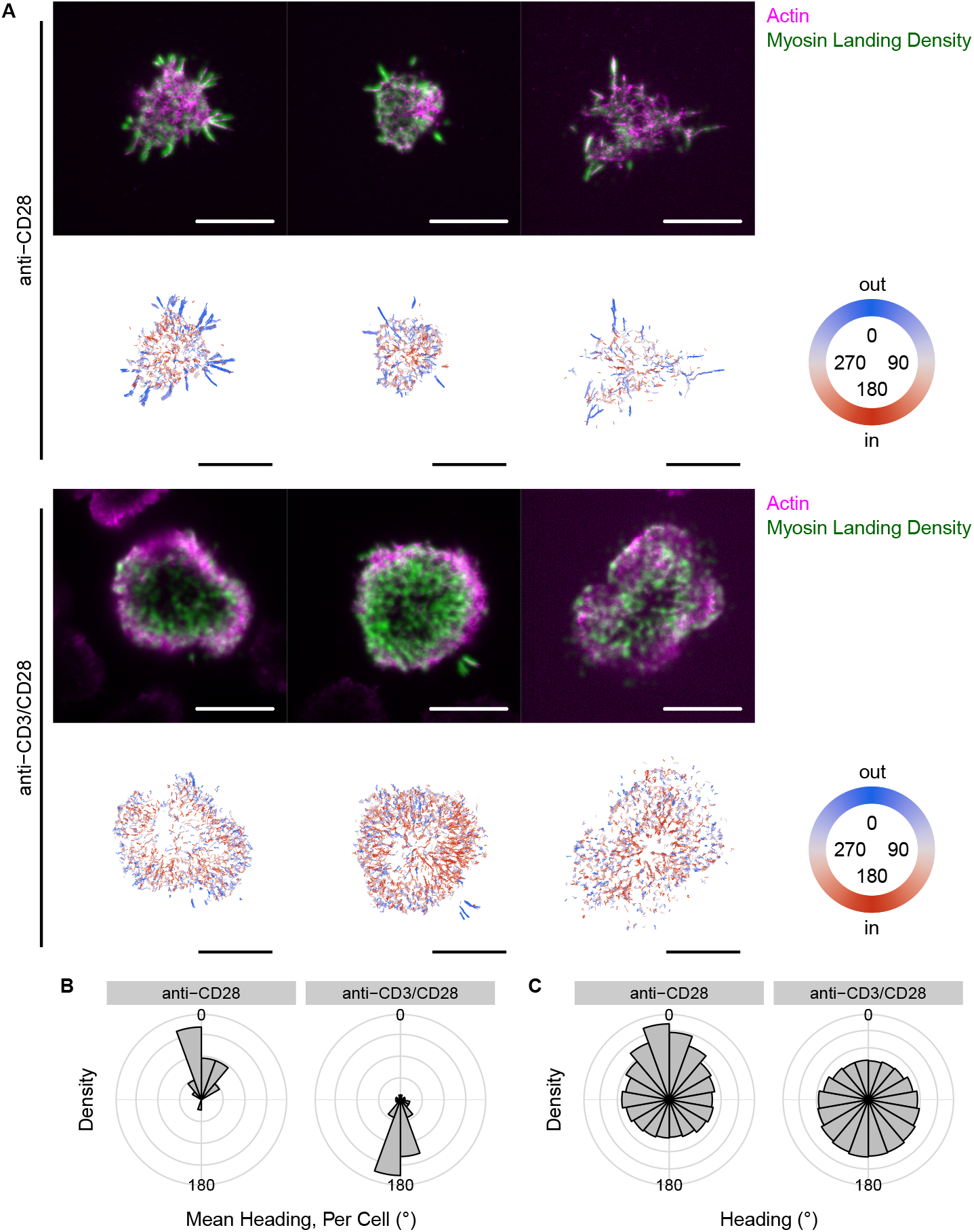
Myosin-5 reverses direction, heading inward at the immunological synapse. (A) Representative Jurkat T cells in contact with non-activating (anti-CD28) or activating (anti-CD3/CD28) surfaces. The top row of each set shows actin (TMR-phalloidin staining) and a kernel density estimate of the myosin-5 landing events. The bottom row in each set shows the detected runs, colored by heading. Headings of 0° run out from the cell center, while headings of 180° run in toward the cell center. Rose plots show the distributions (PDFs) of mean headings on a cell-by-cell basis (B), and headings for all runs collectively (C). Both measures show the reversal of myosin-5 headings on the activating surfaces. All four distributions in (B, C) are nonuniform (Watson test of uniformity; U^2^ = 0.73, 0.82, 13, 3.3; p < 0.01 for each). The heading distributions also differ within (B) and within (C) (Watson-Wheeler test of homogeneity; W = 27, 570; p = 9 × 10^−7^, < 2 × 10^−16^). Scale bars, 10 μm.

On the activating anti-CD3/CD28 surfaces, the Jurkat cells form synapses. Over several minutes, new actin polymerizes at the distal edge to fill out the space between microvilli with a dense network (the dSMAC / pSMAC, see Fig. 1). We find anomalous inward-directed myosin-5 motility on these synapses (Fig. 2A-C and Video S2). This inward heading is inconsistent with the known barbed-end-out orientation of the Arp2/3 nucleated networks at the edge of the IS. Indeed, our own recordings of actin filament polymerization upon IS formation show the stereotypical pattern of barbed-end-out actin nucleation and retrograde flow at the outer edge (Video S3) [47]. However, most inward myosin-5 runs occur in the hypodense actin network at the center of the synapse (the cS-MAC), rather than in the outer and more dense lamellopodial actin. Myosin-5 selects this low-density central network [47], which has a majority of filaments with a barbed-end-in orientation as indicated by the direction of the myosin-5 runs. Presynaptic cells have net-outward myosin-5 traffic, while synaptic cells have net-inward myosin-5 traffic (Fig. 2B).

### The reverse gear myosin-6 motor also reroutes at the synapse

Myosin-6 is a second class of processive myosins that travels toward the pointed end of actin filaments, unlike myosin-5 and all other characterized myosins [44, 35]. This functional difference led us to believe that myosin-6 might differ in its actin network preferences and motility at the IS, which motivated similar myosin-6 tracking experiments. On the anti-CD28 surfaces myosin-6 tends to move inward, as seen in the predominantly red runs in Fig. 3A (see also Video S4). Microvilli support frequent inward myosin-6 runs, again consistent with the known polarity of these actin-based protrusions.

**Figure 3:**
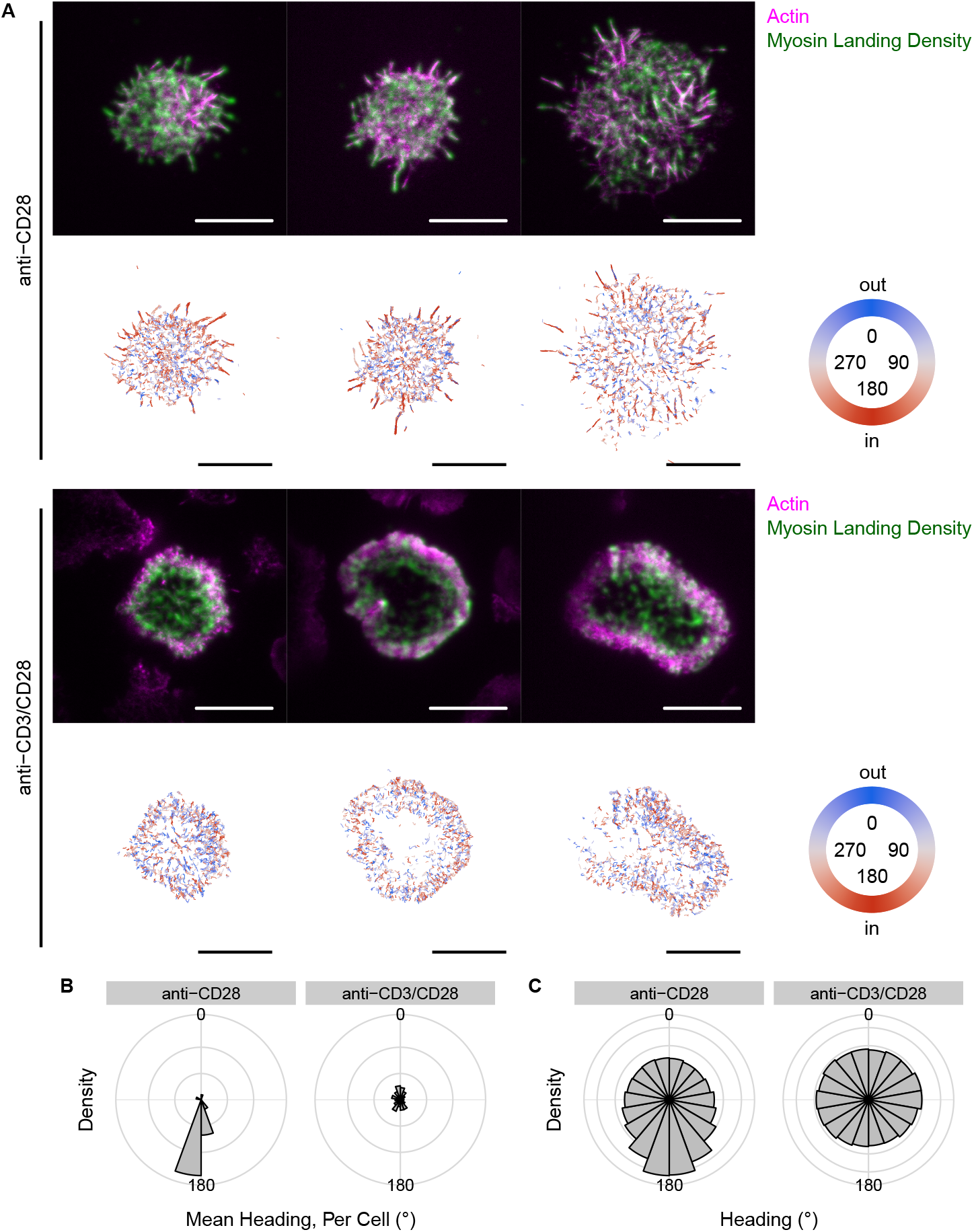
Myosin-6 transitions from inward to nearly isotropic motility upon synapse formation. (A) Representative Jurkat T cells in contact with non-activating (anti-CD28) or activating (anti-CD3/CD28) surfaces, with myosin-6 landing density and runs displayed as in Fig. 2. Rose plots show the distributions (PDFs) of mean headings on a cell-by-cell basis (B), and all run headings combined for all cells (C). Both measures show the inward myosin-6 motility converting to nearly isotropic motility on the activating surfaces. Nevertheless, all four distributions in (B, C) are nonuniform (Watson test of uniformity; U^2^ = 1.9, 1.6, 27, 4.0; p < 0.01 for each). The heading distributions also differ within (B) and within (C) (Watson-Wheeler test of homogeneity; W = 35, 750; p = 3 × 10^−8^, < 2 × 10^−16^). Scale bars, 10 μm.

On the anti-CD3/CD28 surfaces, myosin-6 travels nearly isotropically (Fig. 3A and Video S5). The mean cellular heading distribution has two balanced and less prominent lobes (Fig. 3B). The heading distribution for all runs (Fig. 3C) is also nearly isotropic, although with a preference for circulation (ellipsoidal distribution with maxima at 90° / 270°). The observation that the myosin-6 headings do not mirror those of myosin-5 is evidence that these two myosins select different actin networks that differ in orientation. Myosin-6 lands more evenly throughout the T cell compared to myosin-5, in particular on the lamellopodial actin of the IS.

### Myosin headings depend upon the radial location within the cell

The immunological synapse constructs three zones of signaling molecules called the supramolecular activation clusters, which vary in actin network architecture (Fig. 1, bottom) [18]. The outermost zone (the dSMAC) has Arp2/3 dendritic actin oriented with their barbed ends out. The intermediate zone (the pSMAC) has circumferential bands of actin that bundle and contract under nonmuscle myosin-2 activity. Finally, the central zone (the cSMAC) is hypodense and poorly characterized. Because the actin orientations vary in these three zones, we next examined whether we could detect myosin rerouting across the zones.

Upon synapse formation, both myosin-5 and myosin-6 landing events shift out from the cell center but to a differing extent (Fig. 4A). One significant contributor to this shift is the nucleation of new actin between microvilli, which increases the accessible area for myosin runs at the outer extent of the cell. With anti-CD28 surfaces, myosin-6 runs concentrate in the interior of the cell. This concentration may reflect myosin-6’s preference for older actin at the interior of the cell [48].

**Figure 4:**
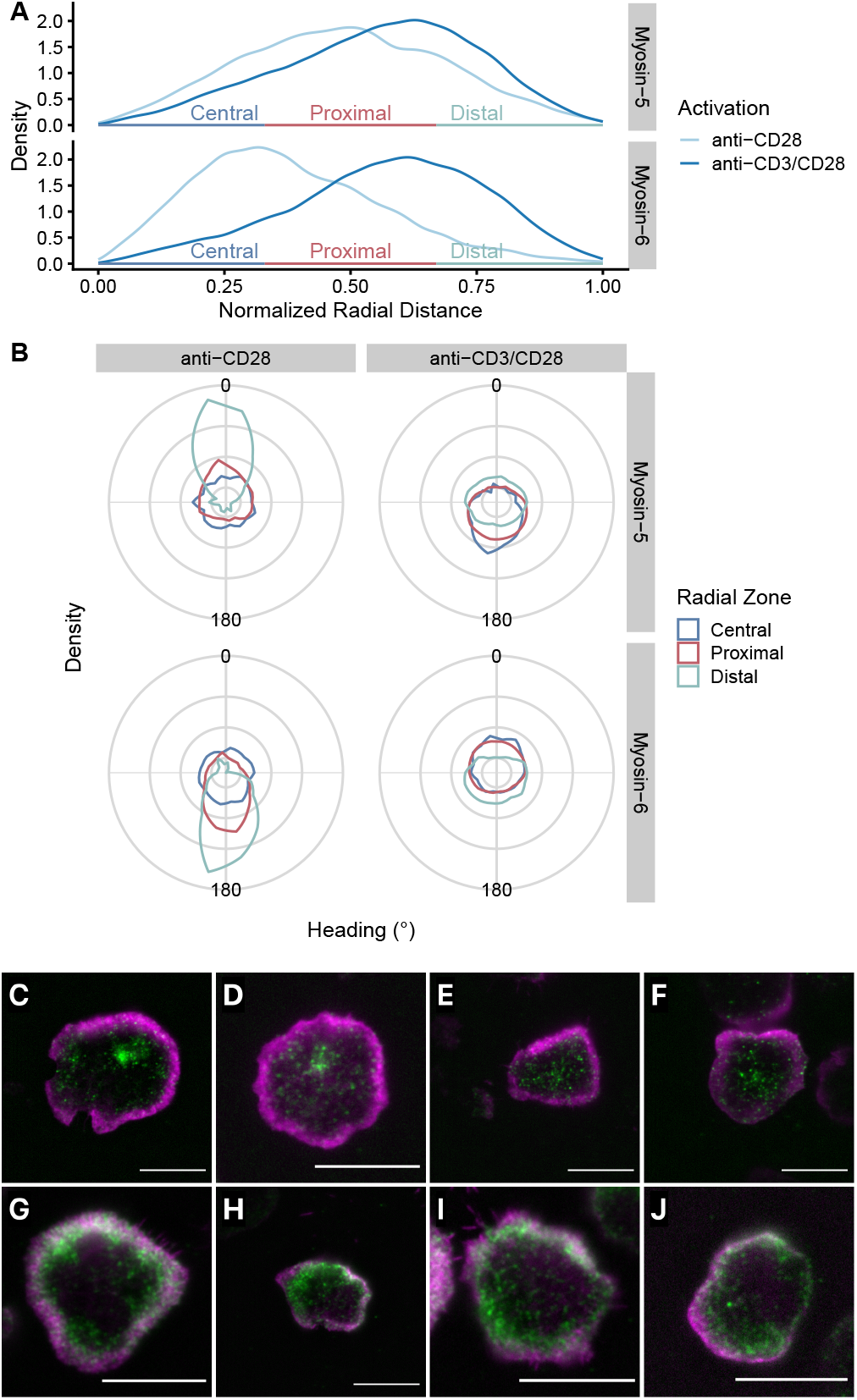
Myosin motility depends upon the radial location within the T cell. (A) Distributions of radial distances from the cell center to the start of myosin runs. To allow comparisons between cells, we normalize distances relative to the maximum radial distance observed for each cell. On activating anti-CD3/CD28 surfaces, runs of both myosin-5 and myosin-6 shift outward. We divide the radial range into thirds, defining central, proximal, and distal zones that approximate the cSMAC, pSMAC, and dSMAC. (B) Heading changes depend upon the radial zone. On the non-activating anti-CD28 surfaces, both myosin-5 and myosin-6 show strong outward and inward preferences (respectively) in the distal zone. On activating anti-CD3/CD28 surfaces, myosin-5 moves nearly isotropically through the distal zone, but moves inward in the central and proximal zones. Likewise, myosin-6 moves outward from the central and proximal zones, and inward from the distal zone. This motility pattern would tend to focus myosin-6 at the proximal-distal boundary. The three zone distributions differ within each of the four conditions (Watson-Wheeler test of homogeneity; W = 600, 740, 550, 800; p < 2 × 10^−16^). Immunofluorescence images show that endogenous myosin-5 concentrates at the cSMAC in either a tight central cluster (C, D), or in a more scattered central pattern (E, F). Endogenous myosin-6 concentrates at the pSMAC/dSMAC (G-J). Magenta: actin, green: myosin, scale bars, 10 μm.

To distinguish myosin motility across the SMACs, we divided the myosin runs into three radial zones—central, proximal, and distal—and investigated the heading distributions within each of the three (Fig. 4B). On the anti-CD28 surfaces, both myosins are strongly polarized (5:out and 6:in) in the distal zone containing the microvilli. That polarization decays to isotropic motility through the proximal and into the central zones. On the anti-CD3/CD28 surfaces, myosin-5 moves inward in the central and proximal zones, and move nearly isotropically but with an outward bias in the distal zone. In contrast, myosin-6 moves out from the inner two zones and in from the outer zone. Thus, myosin-5 motility diverges at the proximal / distal interface, while myosin-6 converges.

Based upon these headings over radial zones, we predict that myosin-5 concentrates in the cSMAC, while myosin-6 concentrates at the pSMAC / dSMAC boundary. We used immunofluorescence microscopy to determine if endogenous myosins localize in these predicted locations. Myosin-5 concentrates either in a narrow region within the cSMAC (Fig. 4C, D), or scatters throughout the cSMAC (Fig. 4E, F). The former, concentrated central pattern resembles the one observed by Bizario and coworkers [5]. Myosin-6 is more abundant, and covers the outermost pSMAC and dSMAC zones (Fig. 4G-J). These localization patterns suggest that both myosins are active and trafficking at the IS.

### The T cell state governs more than just the myosin heading

The rerouting of myosin-5 and myosin-6 traffic is apparent by eye. In fact, we first noticed the rerouting of myosin traffic from inspecting the raw movies. However, other alterations in myosin motility at the IS might be more subtle and would require deeper analysis. Our tracking algorithms allow us to quantify other myosin motility features, including the landing rate, speed, persistence of movement, etc. (see Methods). As we show here, these features provide additional insight into how T cells control their actin networks at the synapse. Because we expect complex correlations between features, we use the UMAP dimension reduction algorithm to cluster cells by their myosin behavior. We then associate these clusters with the cellular state.

Our UMAP embedding for myosin-5 appears in Fig. 5A-C. As expected, the dominant feature is that cells cluster by heading in a circular pattern. Because the embedding orientation is arbitrary, we have rotated and reflected it to align with the color scheme in Fig. 5A. The anti-CD28 and anti-CD3/CD28 Jurkats cluster into two groups at opposite ends (Fig. 5B), flanking a central ring-like cluster (see below). The anti-CD28 cells cluster at the top (0°) while the anti-CD3/CD28 cells appear at the bottom (180°).

**Figure 5:**
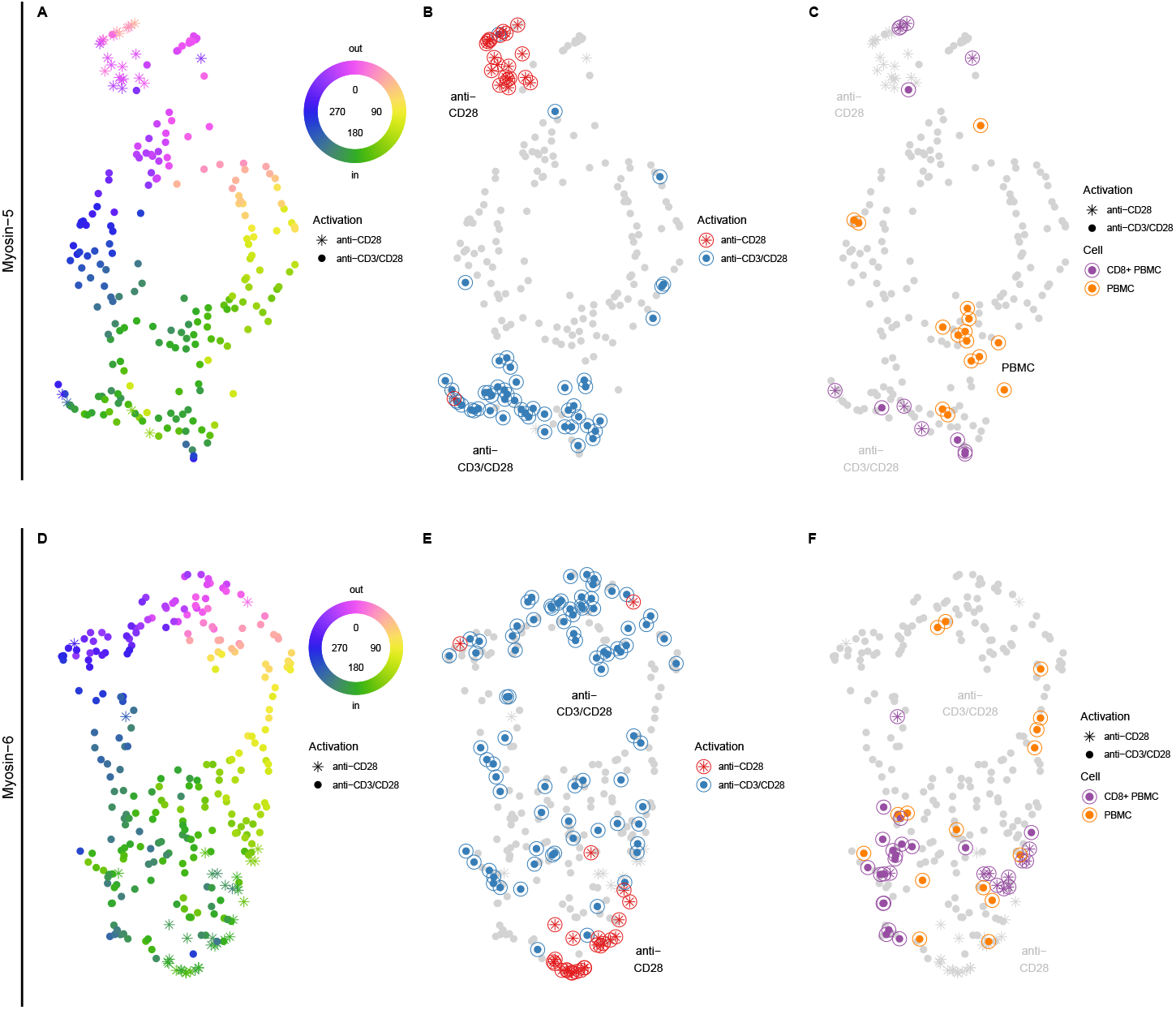
Myosin motility reports the T cell state. We applied UMAP dimension reduction to visualize clusters of myosin behavior, using feature vectors derived from all myosin run properties (see methods). We analyzed myosin-5 and myosin-6 treated cells as separate groups. (A) UMAP embedding of myosin-5 treated cells (including all cell lines, conditions, and treatments), colored by the median heading of the cell. The embedding has ring shaped central cluster with lobes extending out at 0° and 180°. Point shapes that mimic the shape of T cells indicate the surface conditions. (B) Highlighted points show Jurkat cells activated with anti-CD3/CD28 near 180°, or via CD28 alone near 0°. These groups form the two extended lobes. Text labels annotate the embedding regions, while the red/blue color indicates the actual coverslip condition for the individual cells. (C) Highlighted points show PBMCs and CD8^+^ T cells clustering near their Jurkat counterparts. (D–F) The same as in (A–C), but for myosin-6. Note the loose cluster of anti-CD3/CD28-stimulated cells scattered around the central ring, reflecting the more isotropic motility of this myosin at the synapse.

Because Jurkat T cells are an immortalized line with potential irregularities that might affect the construction of actin networks [1, 14], we also examined human PBMCs in our motility assay. The PBMCs on anti-CD3/CD28 surfaces and probed with myosin-5 (Fig. 5C) appear near the corresponding Jurkats but offset to the upper-right. These cells show the same heading reversal as the Jurkat T cells, as seen in the green color of these points in Fig. 5A. However, other features differ somewhat (Supplementary Fig. S1, S2), allowing the PBMCs to form their own cluster here. Because Jurkat T cells have a CD4^+^ origin, we also purified by negative selection CD8^+^ PBMCs to understand a second major T cell class. These CD8^+^ cells cluster within the Jurkat anti-CD3/CD28 cluster, suggesting they share similar motility features. Thus, the myosin traffic rerouting we see at the immune synapse appears to be a general phenomenon.

The UMAP embedding of cells probed with myosin-6 shows a similar circular feature, but with less distinct outer clusters (Fig. 5D-F). The Jurkat cells on anti-CD28 coverslips appear at the bottom of the cluster and are inward directed. However, the synapse-forming cells on anti-CD3/CD28 coverslips are distributed around the ring, as expected from the rose plots in Fig. 3B-C.

### Actin network inhibitors reroute myosin traffic and affect the generation and display of T cell activation markers

Rerouting at the IS shows that T cells exert control over their actin networks and myosin motors, directing myosins toward minor populations of filaments for their motility. We wondered about the impact of disrupting these myosin traffic patterns on T cell function. To disrupt the actin networks we applied two compounds, CK666 and TR100. The small molecule CK666 inhibits Arp2/3 actin nucleation [30] and thus affects the lamellopodial actin at the dSMAC [29]. TR100 inhibits a specific tropomyosin splice isoform, Tpm3.1 [40]. Because Tpm3.1 directs myosin-5 and nonmuscle myosin-2 activity [37, 3], we expected it to be an informative target.

Jurkat T cells treated with CK666 on anti-CD3/CD28 surfaces construct an atypical IS with many spikes at the distal edge. These synapses appear the same as those in prior work from the Hammer group [29]. They showed that treatment with CK666 causes the collapse of the lamellopodial actin, leaving behind formin-nucleated actin filament bundles (Fig. 6A). These long and linear bundles are enhanced by the increased pool of available G-actin [9]. Both myosin-5 and myosin-6 land and move unidirectionally along these bundles. Because they are formin nucleated, the actin bundles have a barbed end out orientation. Therefore, the myosin headings revert to their presynaptic, anti-CD28 preferences (Fig. 6B, compare to Fig. 2–3).

**Figure 6:**
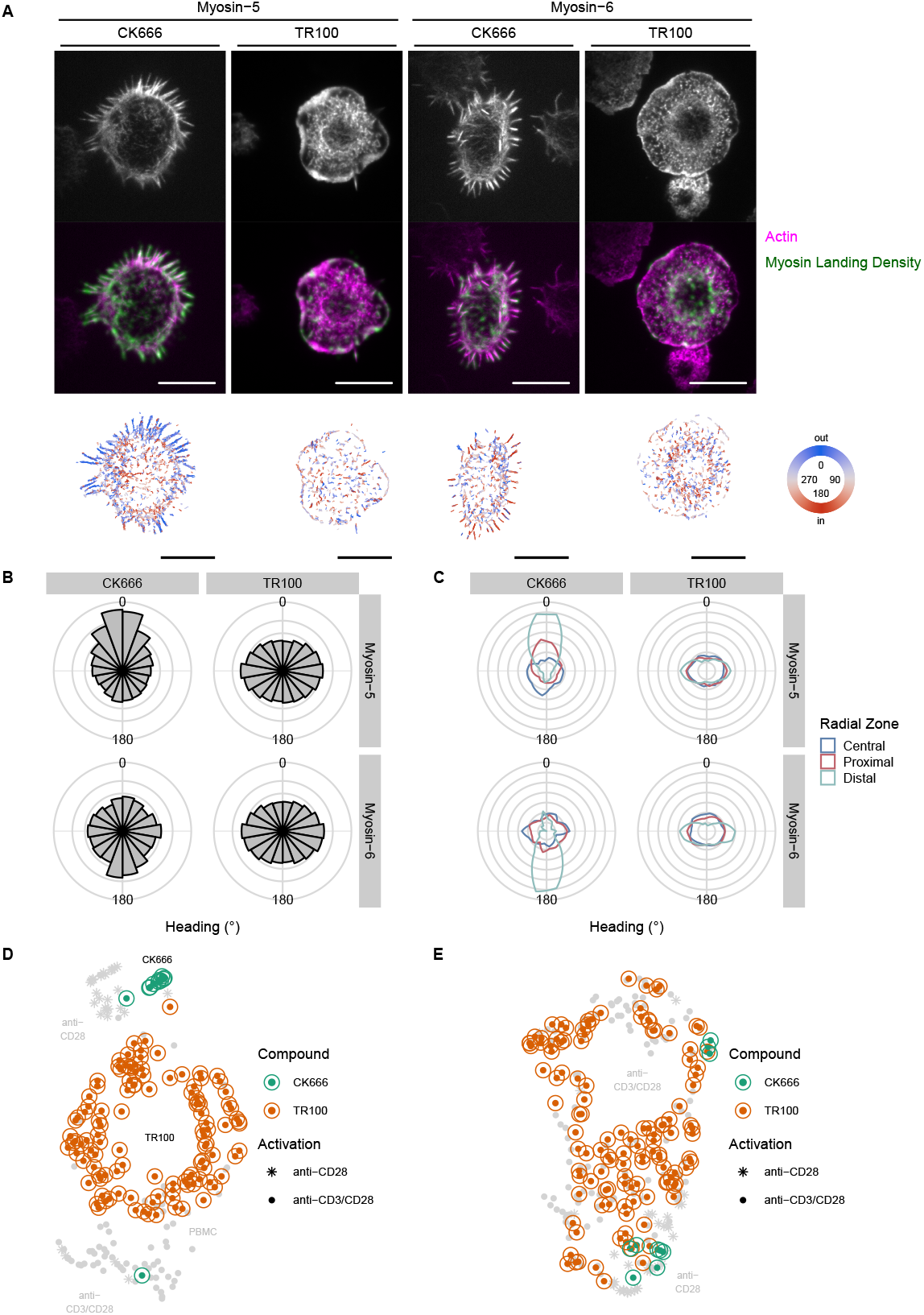
Cytoskeletal inhibitors reroute myosin traffic in T cells on activating surfaces. We treated Jurkat T cells with an Arp2/3 inhibitor (CK666) or a tropomyosin inhibitor (TR100) and applied them to anti-CD3/CD28 surfaces. (A) Fluorescence micrographs of actin, and an overlay with myosin landing intensity displayed as in Figs. 2-3. CK666 treated cells show peripheral spikes that resemble the microvilli of untreated Jurkats on anti-CD28 surfaces. TR100 cells have a similar round morphology to the untreated cells shown earlier, but with differences in the radial distribution of actin. The actin in these TR100-treated cells concentrates in a thin distal zone, with a more diffuse interior before fading in the hypodense central zone. The bottoms row shows myosin runs colored by heading. Scale bars, 10 μm. (B) Heading rose plots, of all runs as in Figs. 2-3. CK666 treatment restores the outward myosin-5 traffic and the inward myosin-6 traffic. TR100 treatment causes both myosins to circulate clockwise and counterclockwise. All four distributions are nonuniform (Watson test of uniformity; U^2^ = 36, 3.0, 2.0, 5.9; p < 0.01 for each). (C) Heading distributions of all runs by radial zone, as in Fig. 4. (D) Myosin-5 and (E) myosin-6 UMAP embeddings, as in Fig. 5 but colored to indicate the inhibitor treatment. CK666-treated cells tend to cluster with the nonactivated, anti-CD28 cells. TR100-treated cells form a large ring cluster that included cells that sample all possible headings (compare with Fig. 5A, D).

When treated with TR100, the Jurkat synapses retain their round shape with a smooth outer boundary. However, a close examination of the actin reveals differences in its organization. The TR100-treated cells often show a thin distal ring of dense actin, a broad and diffuse proximal zone, and a central hypodense region. Myosin-5 and myosin-6 both circulate at the outer edge of the synapse. Circulation is indicated by the desaturated colors of the outermost myosin runs in Fig. 6A. Circulation is also found in the ellipsoidal heading rose plots of Fig. 6B, with peaks at 90° and 270°.

When examining the runs separated into radial zones, both compounds strongly affect the headings in the distal zone (Fig. 6C). Although CK666 reroutes myosin-5 out at the distal and proximal IS, central IS traffic is still inward (Watson test of uniformity, U^2^ = 1.0, p < 0.01). Thus, unlike TR100, CK666 does not appear to affect the construction of actin networks at the cSMAC.

The myosin-5 UMAP embedding shows quite distinct clusters for the CK666- and TR100-treated cells (Fig. 6D). The CK666-treated cells appear in the top cluster along with the untreated anti-CD28 cells. The TR100 cells fill out the central ring-shaped cluster because they lack a single directional bias (clockwise and counterclockwise runs are balanced). The myosin-6 UMAP embedding shows similar features (Fig. 6E), except that the TR100 cells are distributed within the untreated anti-CD3/CD28 cell cluster because both sets lack a directional bias.

Given the strong effects of CK666 and TR100 on the immunological synapse, we expected that these compounds might impair other aspects of downstream signaling in T cell activation. To uncover any impact on downstream events, we examined how these two compounds affect the display of activation markers on the T cell surface by flow cytometry. With or without CK666 inhibition, the anti-CD3/CD28 activated Jurkat T cells showed the same levels of CD69 and CD25 activation markers on the cell surface (Fig. 7 and Supplementary Fig. S3). Therefore, even though dSMAC actin is prominent at the IS, its specific disruption is relatively unimportant for downstream T cell activation processes. In contrast, treatment with TR100 significantly reduces surface expression of both CD69 and CD25. Even higher TR100 concentration and extended exposure is cytotoxic to the T cells (Supplementary Figs. S4 & S5). To determine if this reduction in surface markers is due to a trafficking defect, we examined the expression levels of the early activation marker (CD69) in fixed and permeabilized cells. We see a similar reduction in CD69 levels, suggesting that TR100 affects CD69 expression rather than later trafficking events (Supplementary Fig. S6).

**Figure 7:**
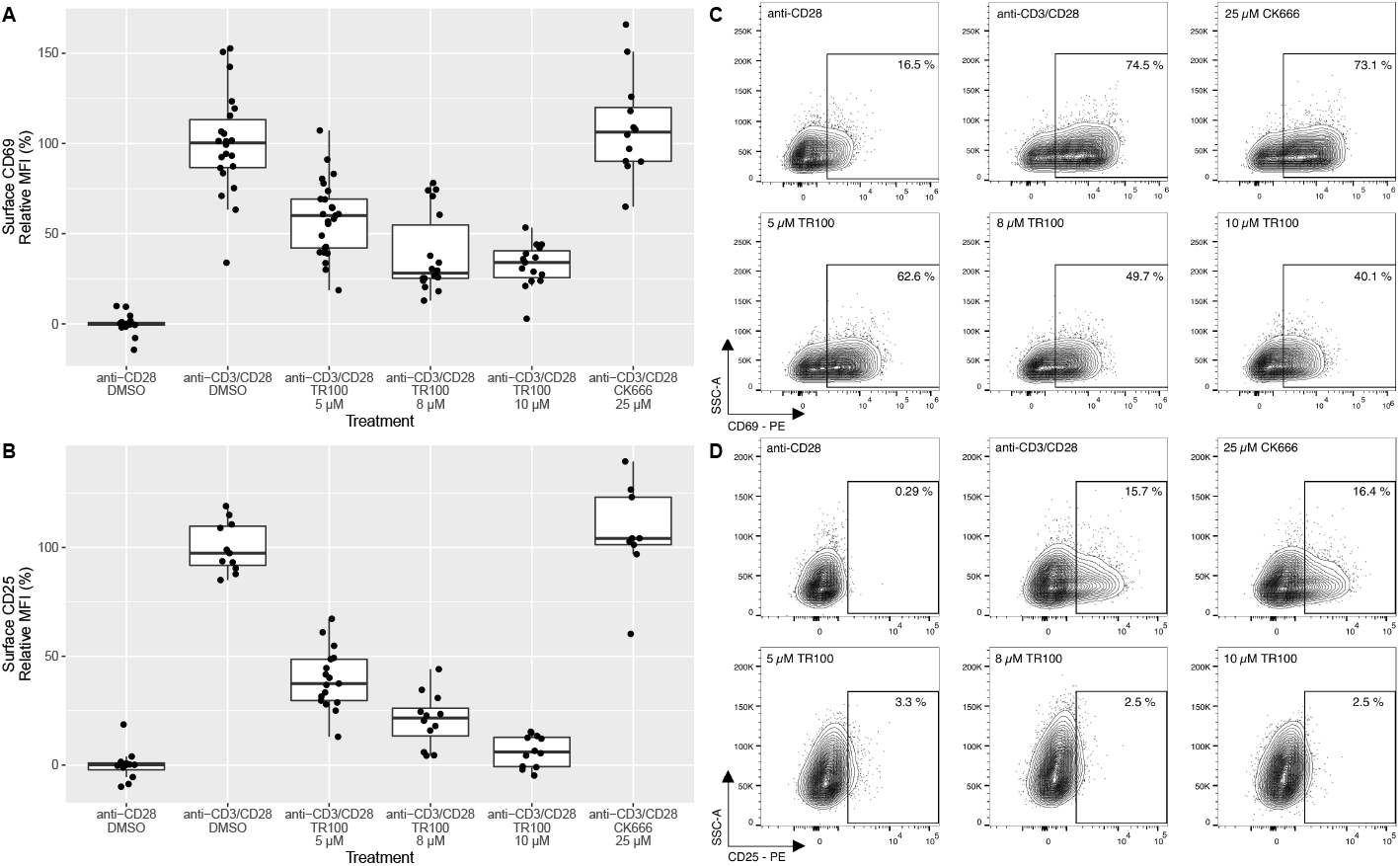
Treatment with TR100 inhibits T cell activation marker display, while CK666 shows no such effect. Flow cytometric analysis reveals a TR100 dose-dependent decrease of (A) surface CD69 (χ^2^(4) = 36.6, p = 2.2 × 10^−7^), and (B) surface CD25 (χ^2^(4) = 35.5, p = 3.6 × 10^−7^), in Jurkat T cells activated with anti-CD3/CD28. Treatment with CK666 does not alter any of these marker levels. Representative plots of (C) CD69 and (D) CD25 display changes after inhibitor treatment. Data points in (A, B) are from N = 3–6 biological replicates with 2–3 technical replicates for each. The median fluorescence intensity (MFI) values are background-subtracted using the mean anti-CD28 values and normalized to the anti-CD3/CD28 control values for each biological replicate. We measured CD69 after 4 h and CD25 after 24 h.

## Discussion

In this work, we have tracked the motion of unconventional myosin motors over the actin networks of the immunological synapse. By tracking these myosins, we construct a functional map of the synapse. This map captures spatial information on the actin network structure, filament polarity, density, and intrinsic myosin preferences for distinct actin networks. The key finding is an aberrant myosin motility after synapse formation, such that myosins move in the opposite direction than one would predict from the actin polymerization patterns. Moreover, the T cell state tunes the motile properties of myosin motors, which allows us to determine the T cell state solely from the information contained within the myosin trajectories. Specific disruptions of actin filament networks affect the display of T cell activation markers, establishing functional linkage between actin networks and T cell activity.

Our understanding of how processive myosin motors navigate the cell is still developing. However, our field has proposed several molecular mechanisms for actin networks to regulate myosin traffic [36]. Actin filament age, applied forces (tension, compression, or torsion), posttranslational modifications, decoration with actin-binding proteins, and higher-order organization in crossings or bundles may all contribute to myosin regulation. Each of these actin network features modulate the interaction interface with myosin motors, which could enhance or inhibit motility in a myosin class-specific manner. Although the IS can control myosin traffic in many ways, our results identify tropomyosins as a key regulator given the impact of TR100 inhibition. Tropomyosins can cooperatively decorate filaments, making them ideal to mark specialized, long actin filaments that direct myosin-5 toward the cSMAC. Tropomyosins regulate myosin-5 traffic in several known and characterized systems [22, 37, 3]. Our TR100 results also identify tropomyosins as a potential molecular target in immune systems. Because inhibition of tropomyosins suppresses CD69 and CD25 production, TR100 may have immunosuppressive activity in addition to its proposed anti-metastatic activity [15, 6, 40].

The *ex vivo* traffic maps reveal a surprising degree of myosin traffic at the cSMAC, given its low F-actin density. The cSMAC is often characterized as actin depleted, hypodense, or devoid of actin [18]. Indeed, one of the proposed roles for actin at the cSMAC is that it inhibits secretion of cytotoxic granules [33, 34, 12]. In this picture, actin at the cSMAC serves as a physical barrier that the T cell must clear for secretion. Our results assign an added role to cSMAC actin, to focus myosin-5 traffic toward the center of the cSMAC. Although it is tempting to speculate that myosin-5 transports TCR microclusters to the cSMAC, it seems clear that other mechanisms suffice. These include frictional coupling of TCRs to actin retrograde flow in the dSMAC and pSMAC, followed by handoff to dyneins in the final phase at the cSMAC [47]. Separate myosin-5 cargoes, including those coupled through various Rab family proteins [27, 45, 32], may instead play a role in T cell activation and IS development.

Our traffic maps also reveal that myosin-6 concentrates at the pSMAC, traveling in from the outside, out from the inside, and circulating in the middle. The pSMAC forms an adhesive ring with the target cell, typically through the combination of LFA-1 and ICAM1 [29]. Although myosin-6 is not known to interact directly with LFA-1, it may have related roles that require localization at the pSMAC. These roles include endocytosis [11, 16, 39] or the anchoring of lipid membranes [21, 2]. Our imaging of endogenous myosins shows that myosin-6 is far more abundant than myosin-5 in T cells. This composition makes sense given that myosin-6 covers a larger area than the small central disk of myosin-5. The T cell may regulate expression levels of these two myosins to ensure coverage of their final target zones.

This work raises several important questions. Although it seems likely that the trafficking actin network is important for IS function given the impact of TR100 inhibition, the exact myosin cargoes are unknown. Such questions are best addressed through proteomics (BioID, APEX, and related methods). Moreover, given the challenges of embedding a trafficking actin network within a greater structural network, we need to determine how the trafficking network assembles. Finally, we must assess the functional diversity of the trafficking actin network across T cell subsets and/or functional states. Are these specialized trafficking networks tuned to support effector mechanisms of T cells? Our data seem to support that idea, but a clearer picture will emerge from work on rarer T cell subpopulations.

Overall, this work has identified a new form of actin network specialization at the immunological synapse. Our results support a model of a network embedded within a network: a minor actin network for myosin trafficking, inside a larger and more prominent actin network to set the synaptic shape. These findings motivate future work to understand the structural dynamics and functional adaptations of the actin networks at the synapse.

## Acknowledgments

We thank John Hammer for sharing unpublished data and for the tdTomato-F-Tractin construct, Anthony Kossiakoff and members of the Kossiakoff laboratory for assistance with flow cytometry, and Ainhoa Arina for helpful discussions and comments on the manuscript. We acknowledge the University of Chicago Research Computing Center and NIH Grant R01 GM124272 for support of this work.

## Materials and methods

### Cells and reagents

Jurkat cells (clone E6-1) were cultured in RPMI 1640 medium (Thermo Fisher Scientific) supplemented with 10% heat-inactivated FBS (Sigma). We obtained PBMCs derived from anonymous healthy human donors (ZenBio). The CD8^+^ T cells were purified from PBMCs using negative selection bead binding (CD8^+^ T Cell Isolation Kit, human, Miltenyi Biotec). The following compounds were used at the indicated concentrations: TR100 (5, 8 or 10 μM, Sigma Aldrich), CK666 (25 μM, Fisher). Baculovirus containing myosin-5-YFP HMM and myosin-6-GFP HMM expression constructs were used to infect Sf9 cells and the motor proteins were purified using Flag-affinity chromatography as previously described [8, 35].

### Ex vivo motility

Plasma-cleaned glass coverslips were coated with 0.01% poly-L-lysine (Sigma) for 30 min, RT, followed by coating with antibodies: anti-human CD3 (clone OKT3, 10 μg/mL, Invitrogen) and anti-human CD28 (clone CD28.2, 10 μg/mL, BD Pharmingen) for 1 h at 37 °C. Cells were seeded onto prepared coverslips and activated for 5 min at 37 °C. During the inhibitor assays, cells were pre-treated with 1% DMSO as vehicle control or selected inhibitors for 15 min at 37 °C before activation.

Cells were then washed with PEM buffer (0.1 M PIPES, 5 mM EGTA, 2 mM MgCl_2_·6 H_2_O, pH 6.8) containing 3.3% (v/v) polyethylene glycol 8000, 0.083% Triton X-100, 0.83 μg/mL phalloidin-tetramethylrhodamine B isothiocyanate (Sigma), 2.67% paraformalde-hyde, and immediately washed with PBS. Imaging chambers were then assembled using glass coverslips with cells and microscope slides with double stick tape. Imaging chambers were filled with motility buffer (25 mM imidazole·HCl, 1 mM K·EGTA, 4 mM MgCl_2_, 25 mM KCl, pH 7.5) containing recombinant myosins, BSA (1 mg/mL, Calbiochem), 1% Triton X-100, 2 mM ATP, and an oxygen scavenging system composed of 216 μg/mL glucose oxidase (Calbiochem), 36 μg/mL catalase (Sigma), and 4.5 mg/mL glucose.

### Single-molecule imaging using TIRF microscopy

We used a custom-built objective-type total internal reflection fluorescent microscope for single-molecule myosin tracking. The microscope is equipped with a 100x, 1.40 NA objective (Olympus SuperApo) and an EMCCD camera (iXon, Andor Technologies). Movies were collected at 2 Hz. A single frame of tetramethylrhodamine-phalloidin (TMR-phalloidin) stained actin was visualized first, followed by an image stack of the FP to observe myosin movements. Images were collected at 23 °C.

### Myosin tracking

Movies of single-molecule myosin movements were tracked using the TrackMate package in FIJI [13], using previously described procedures and settings [8]. We exported an XML file for each movie that contained the myosin tracks identified by TrackMate. This file was then processed with a python script that calculated a set of motility metrics for each myosin trajectory, as defined here. Persistence is the end-to-end distance divided by the runlength. The persistence is a unitless quantity that approaches one for straight paths, and falls to zero for bent paths or paths with back-and-forth behavior. Heading is the angle between the vector from the start to the end of the path, relative to the vector from the center of the cell to the start of the run. Heading values were unwrapped to fall within the range of 0–360°. Paths that point away from the center of the cell have a heading of 0°, while paths that point toward the cell center have a heading of 180°. The normalized start radius is the distance from the start of the trajectory to the cell center, divided by the maximal distance for each cell. We collected these path metrics for all tracking experiments, along with factors identifying the experimental conditions. Summary statistics for paths were generated in R and plotted using the package *ggplot2*. Landing intensity images in Figs. 2 and 3 were produced using the *spatstat* package (version 2.2) in R.

### UMAP dimension reduction

For each recorded cell, we constructed feature vectors from the five quantiles (10%, 25%, 50%, 75%, and 90%) of the following myosin trajectory metrics: velocity, persistence, normalized start radius, and the x- and y-components of the heading vectors. We also included the landing rate and the circular mean resultant length (rho) as single values for each cell. The mean resultant length is a quantity that expresses the tendancy of a collection of heading vectors from a single cell to point in the same direction. All heading vectors are mapped onto the unit circle, and the arithmetic mean of all points is taken. This mean value will lie within the unit disc, and its radius is assigned to rho. Rho is 0 for headings that are uniformly distributed around the unit circle, and is 1 if all headings point in the same direction. Using this collection of features in our feature vector, we then applied the UMAP algorithm as implemented in the *UMAP*.*jl* package (version 0.1.6) in Julia [4]. The input parameters for UMAP were 50 neighbors, a minimum distance of 0.015, and 50,000 epochs.

### Immunocytochemistry

Jurkat cells were activated following the same protocol as described for *ex vivo* motility. In the next step cells were fixed with 4% paraformaldehyde for 15 min at RT. Cells were then washed with PBS, permeabilized using 0.1% Triton X-100, 3 min at RT, washed with PBS and incubated with blocking buffer (3% BSA in PBS) for 30 min at RT to decrease nonspecific antibody binding. Cells were then incubated overnight at 4 °C with primary antibodies diluted in blocking buffer: anti-myosin-5a antibody (1:100, #3402, Cell Signaling Technology), or anti-myosin-6 antibody (1:200, ABT42, Millipore). Cells were washed with PBS, and incubated for 1 h at RT with Cy5-conjugated goat anti-rabbit antibody (1:250, #A10523, Thermo Fisher Scientific) diluted in blocking buffer. After incubation cells were incubated for 5 min at RT with TMR-phalloidin (0.83 μg/mL) and Hoechst 33342 (1 μg/mL, Invitrogen) in PBS and washed with blocking buffer before imaging on a Zeiss Axiovert 200 equipped with an Andor Luca camera.

### Flow cytometry

Cells treated or untreated with chosen inhibitors were activated at 37 °C for 4 h or 24 h on 96-well plates coated with 0.01% poly-L-lysine and CD28 (10 μg/mL) or CD3/CD28 (10 μg/mL each) antibodies. After activation, cells were stained with antibodies: CD69-PE (clone FN50, Biolegend) and CD25-PE (clone BC96. Biolegend). Selection of live cells was accomplished by staining with either propidium iodide or Fixable Viability Dye eFluor 520 (eBioscience).

Cells were analyzed using a CytoFLEX V0-B2-R2 flow cytometer (Beckman Coulter Life Sciences). Data were analyzed using FlowJo Software (v10.0.7). The obtained median fluorescence intensity (MFI) values were normalized for each separate experiment by transforming the mean MFI of non-activated (CD28 only) cells to 0%, and mean MFI of activated control cells (anti-CD3/CD28 + DMSO) to 100%.

### Data exclusions

We excluded 10 cells after inspection of their heading distributions, 7 probed with myosin-5 and 3 probed with myosin-6. All were Jurkat T cells under activating anti-CD3/CD28 conditions and no cytoskeletal inhibitors. The excluded cells had the characteristic spiky appearance of the unactivated Jurkats shown at the top of Fig. 2A. These cells also showed a preference for outward myosin-5 / inward myosin-6 movements, in contrast with the remainder of the datatset. We suspect that these cells landed on coverslips with defective or incomplete antibody coating.

### Statistical analysis

All significance tests for nonuniform heading distributions were determined using the Watson test of uniformity. Comparisons between heading distributions were determined with the nonparametric Watson-Wheeler test. Both use the implementation found in the R package *circular* (version 0.4-93). For Fig. 7, we used linear mixed effects regression to model the effect of treatment on the CD69 and CD25 cell-surface markers. We used treatment (the three concentrations of TR100 plus the single CK666) as the fixed effect, and the interaction of biological replicate and treatment as random effect intercepts. We used a likelihood ratio test to obtain p-values, comparing to a null model that omits the treatment fixed effect. These linear mixed effects analyses were performed with the R package *lme4* (version 1.1-27.1). For all experiments, α = 0.05 was used as the threshold for statistical significance.

## Supplementary Information

**Table S1:** Summary of experimental conditions, number of cells acquired, and number of myosin runs acquired.

| Motor | Activator | Compound | Cell | N.cells | N.runs |
| --- | --- | --- | --- | --- | --- |
| Myosin-5 | anti-CD28 | Untreated | CD8<sup>+</sup> PBMC | 7 | 776 |
| Myosin-5 | anti-CD28 | Untreated | Jurkat | 21 | 14709 |
| Myosin-5 | anti-CD3/CD28 | CK666 | Jurkat | 11 | 23758 |
| Myosin-5 | anti-CD3/CD28 | Untreated | CD8<sup>+</sup> PBMC | 5 | 1143 |
| Myosin-5 | anti-CD3/CD28 | Untreated | Jurkat | 49 | 65510 |
| Myosin-5 | anti-CD3/CD28 | Untreated | PBMC | 15 | 11284 |
| Myosin-5 | anti-CD3/CD28 | TR100 | Jurkat | 114 | 44454 |
| Myosin-6 | anti-CD28 | Untreated | CD8<sup>+</sup> PBMC | 11 | 3068 |
| Myosin-6 | anti-CD28 | Untreated | Jurkat | 26 | 23840 |
| Myosin-6 | anti-CD3/CD28 | CK666 | Jurkat | 10 | 9967 |
| Myosin-6 | anti-CD3/CD28 | Untreated | CD8<sup>+</sup> PBMC | 18 | 7826 |
| Myosin-6 | anti-CD3/CD28 | Untreated | Jurkat | 87 | 71844 |
| Myosin-6 | anti-CD3/CD28 | Untreated | PBMC | 16 | 5675 |
| Myosin-6 | anti-CD3/CD28 | TR100 | Jurkat | 121 | 55312 |

**Figure S1:**
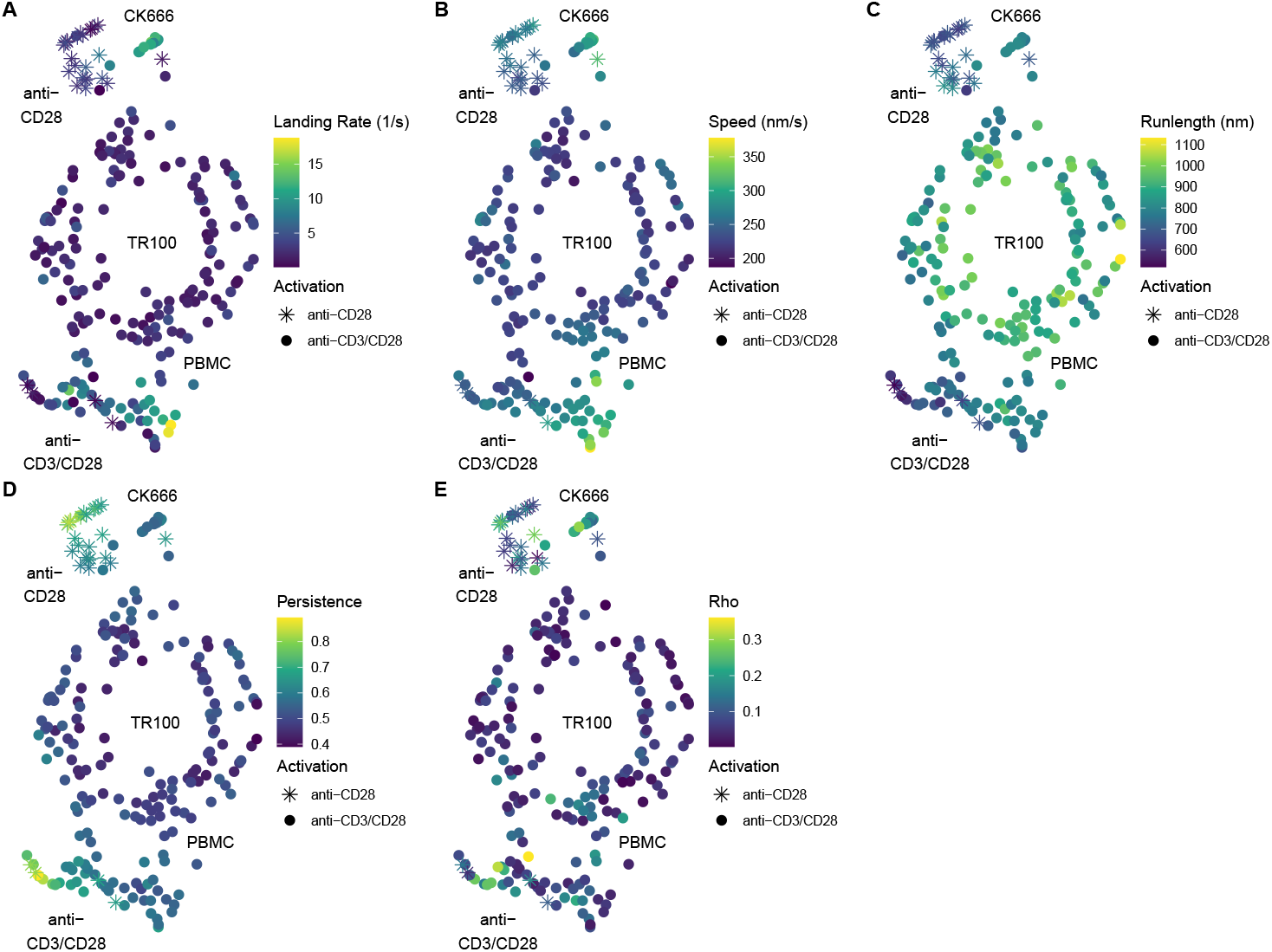
Myosin-5 UMAP embedding features. The points are colored by mean feature values used in constructing the UMAP embedding (aside from the heading, which is shown in 5).

**Figure S2:**
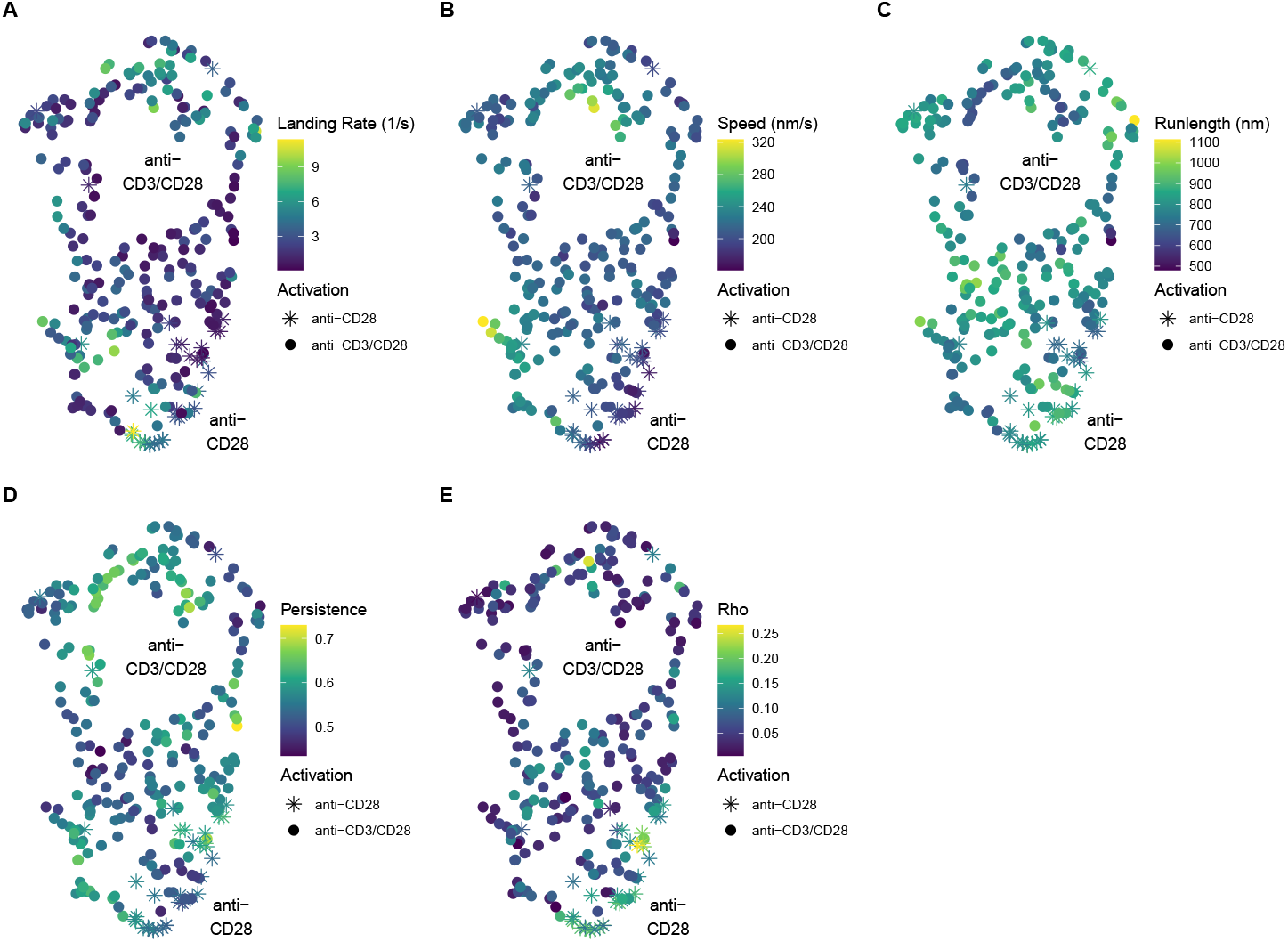
Myosin-6 UMAP embedding features. The points are colored by mean feature values used in constructing the UMAP embedding (aside from the heading, which is shown in 5).

**Figure S3:**
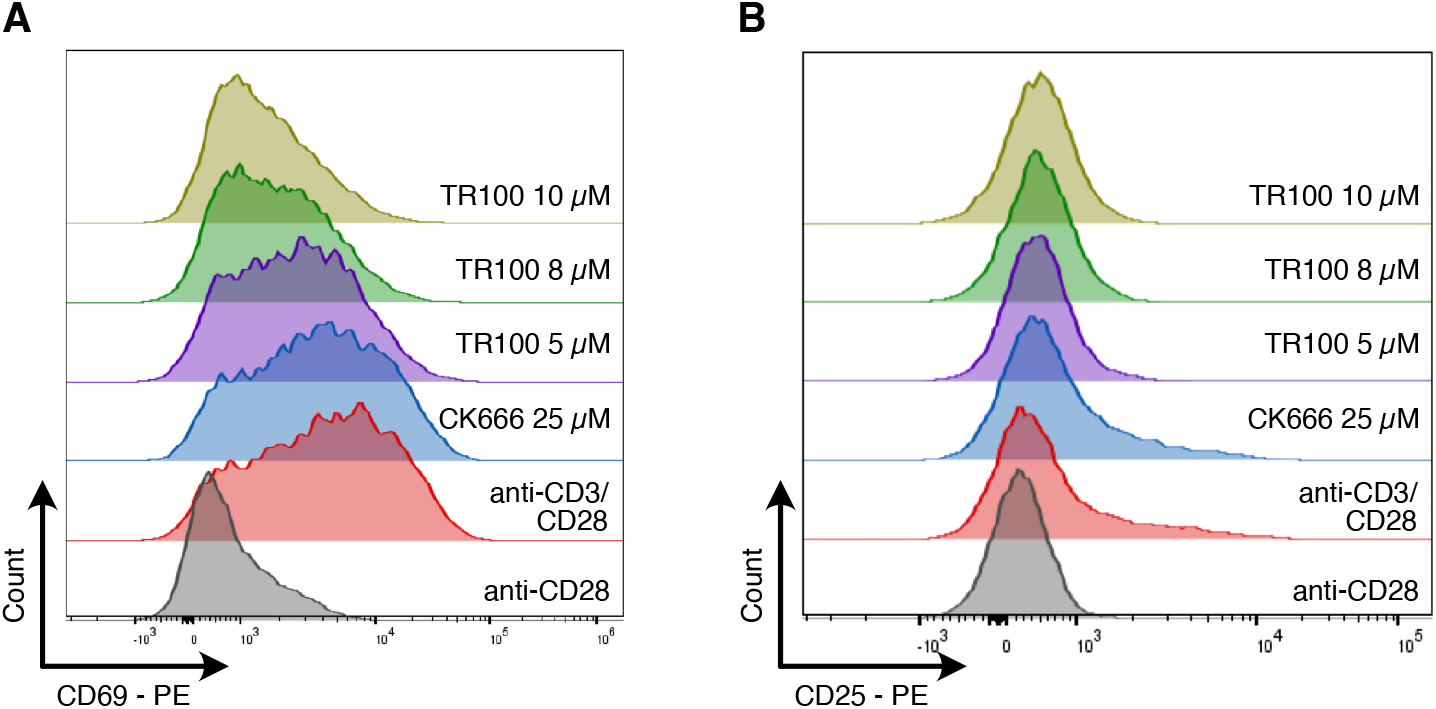
CD69 and CD25 display is inhibited by TR100. Representative distributions of CD69 (A) and CD25 (B) signals measured by flow cytometry. CD69 was measured after 4 h, and CD25 was measured after 24 h.

**Figure S4:**
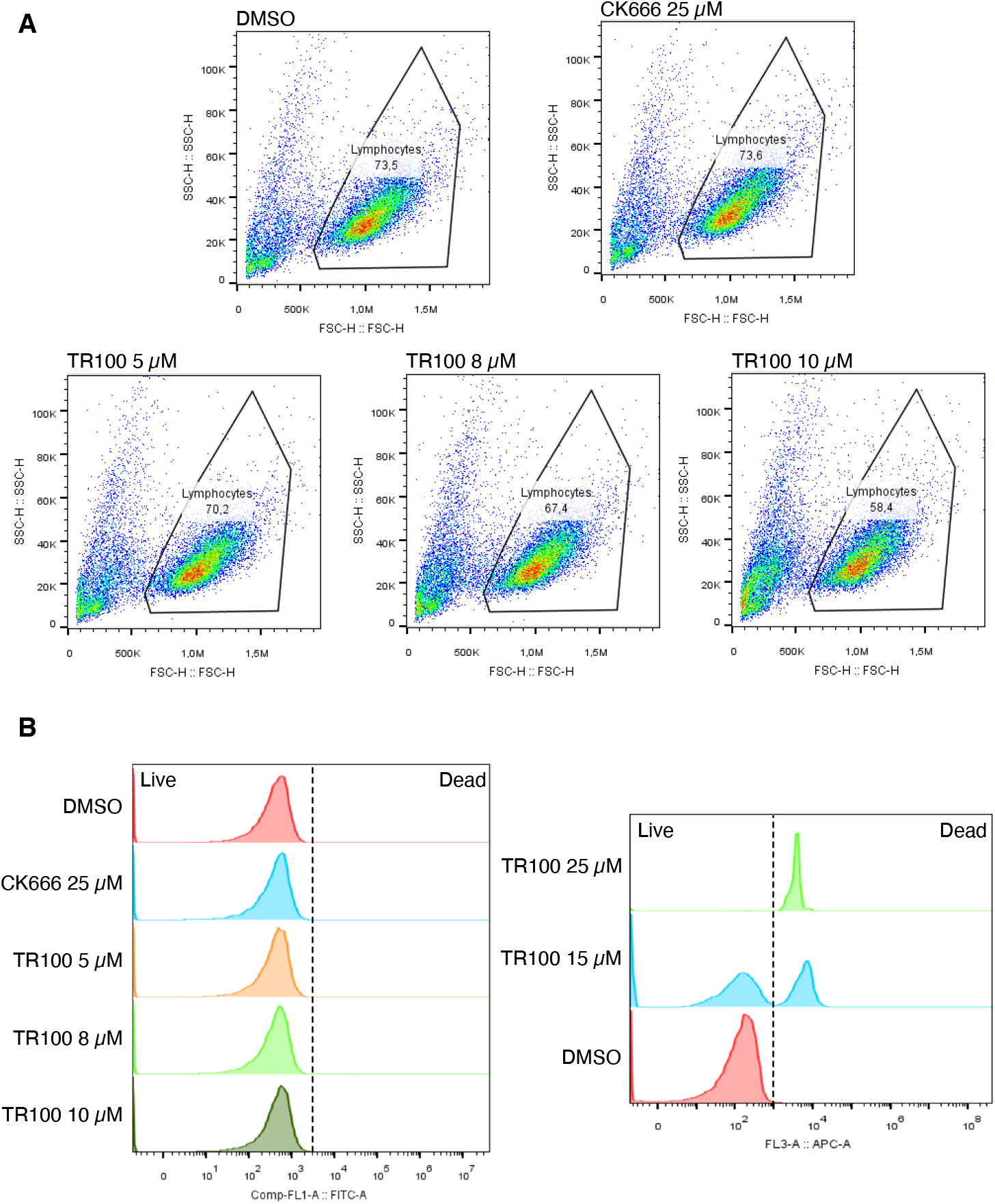
TR100 and CK666 cytotoxicity after 4 h. (A) Representative side scatter / forward scatter (SSC-H/FSC-H) plots for cell inhibitor treatments after 4h (Fig. 7A). (B) Cell viability for inhibitor treatments. Higher concentrations of TR100 (15 μM and 25 μM) result in significant cytotoxicity after 4 h, and were not used in Fig. 7A,C.

**Figure S5:**
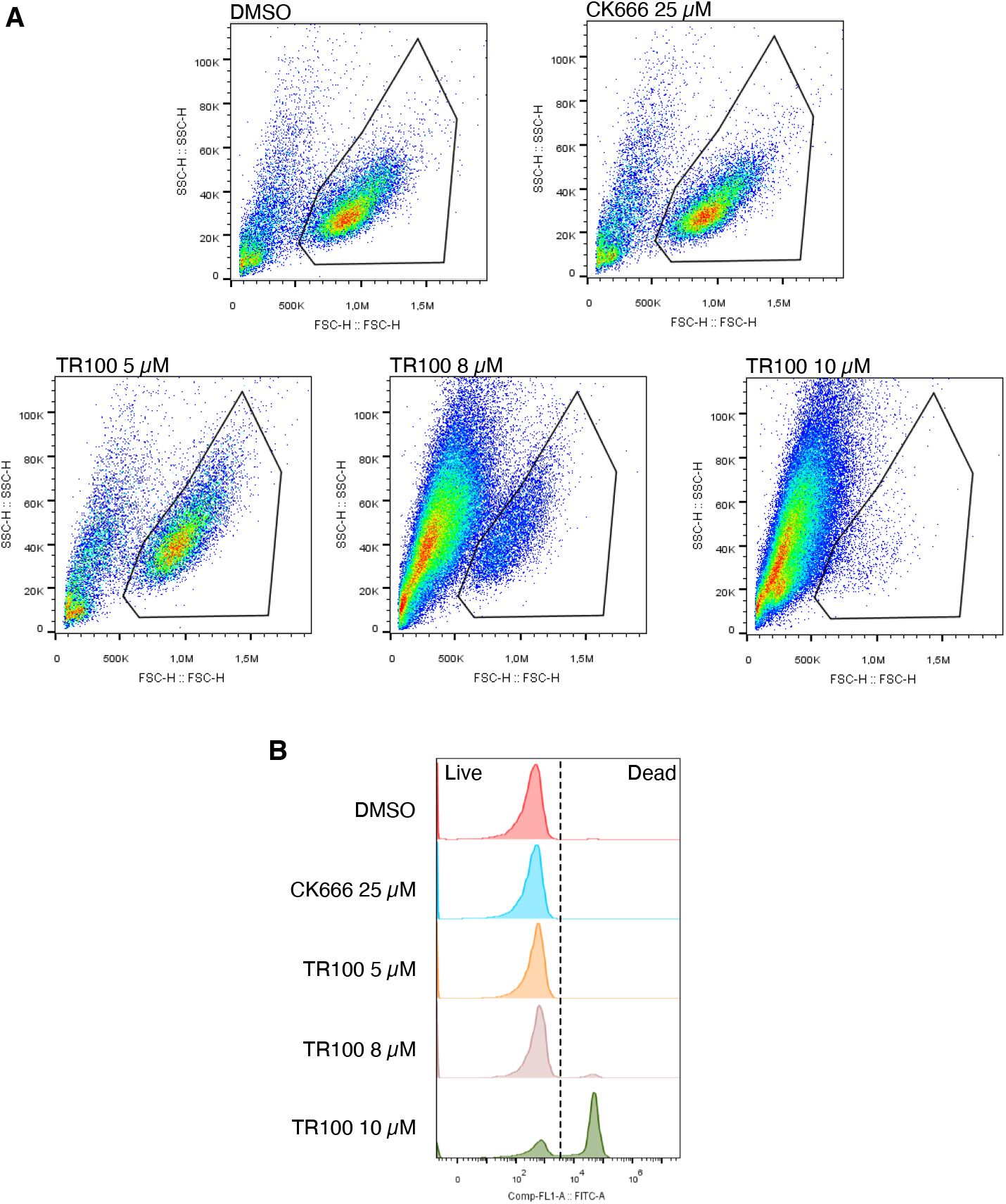
TR100 and CK666 cytotoxicity after 24 h. (A) Representative SSC-H/FSC-H plots for cell inhibitor treatments after 24h (Fig. 7B). (B) Cell viability for inhibitor treatments. The highest concentration of TR100 used here (10 μM) was cytotoxic after 24 h. The remaining viable cells at this TR100 concentration were analyzed in Fig. 7B for CD25. Note that lower TR100 concentrations (5 μM and 8 μM) had sufficient numbers of viable cells for analysis.

**Figure S6:**
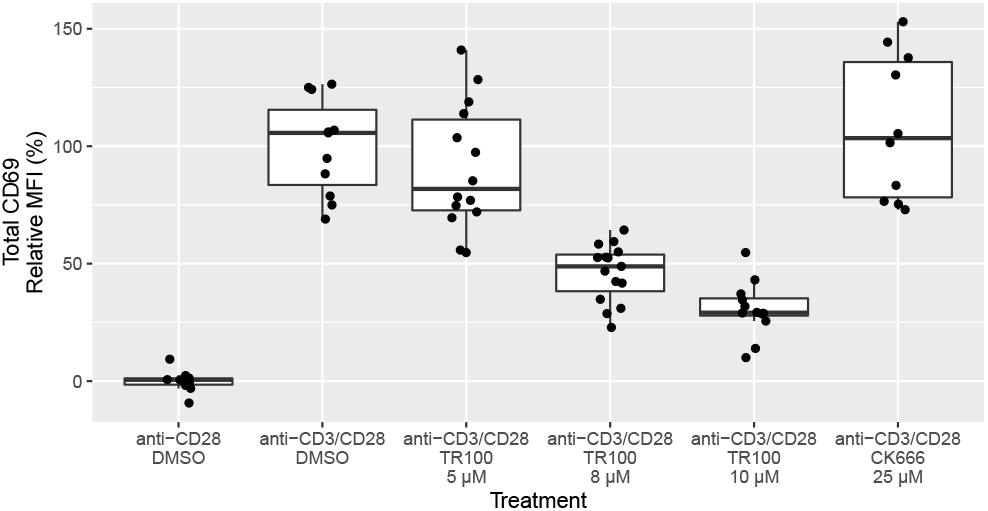
Treatment with TR100 inhibits production of the early T cell activation marker CD69, while CK666 shows no such effect. Flow cytometric analysis of permeabilized cells reveals a TR100 dose-dependent decrease of total CD69 (χ^2^(4) = 53.7, p = 6.1 × 10^−11^), in Jurkat cells activated with anti-CD3/CD28. Treatment with CK666 does not alter CD69 levels. Data points are from N = 3–6 biological replicates with 2–3 technical replicates for each. The median fluorescence intensity (MFI) values are background-subtracted using the mean anti-CD28 values and normalized to the anti-CD3/CD28 control values for each biological replicate. We measured CD69 after 4 h.

*Video S1*

Myosin-5 runs along a Jurkat T cell on an anti-CD28 coverslip. TMR-phalloidin staining of actin in red, myosin in green. The movie shows a 30 × 30 μm area. The movie was recorded at 2 Hz, while playback is at 20 Hz (10x speedup).

*Video S2*

Myosin-5 runs along a Jurkat T cell on an anti-CD3/CD28 coverslip. TMR-phalloidin staining of actin in red, myosin in green. The movie shows a 30 × 30 μm area. The movie was recorded at 2 Hz, while playback is at 20 Hz (10x speedup).

*Video S3*

A Jurkat T cell transfected with tdTomato-F-Tractin, with the video beginning soon after contact with an anti-CD3/CD28 coverslip. Retrograde flow of actin is apparent at the dSMAC once the synapse has maximally expanded. Framerate is 2x live. Scale bar, 5 μm.

*Video S4*

Myosin-6 runs along a Jurkat T cell on an anti-CD28 coverslip. TMR-phalloidin staining of actin in red, myosin in green. The movie shows a 30 × 30 μm area. The movie was recorded at 2 Hz, while playback is at 20 Hz (10x speedup).

*Video S5*

Myosin-6 runs along a Jurkat T cell on an anti-CD3/CD28 coverslip. TMR-phalloidin staining of actin in red, myosin in green. The movie shows a 30 × 30 μm area. The movie was recorded at 2 Hz, while playback is at 20 Hz (10x speedup).

